# Improved Stability and Interpretability of Motor Modules Computed with an Autoencoder

**DOI:** 10.1101/2025.11.11.687901

**Authors:** Siddharth R. Nathella, Bryant A. Seamon, Aaron J. Young, Lena H. Ting

**Affiliations:** George W. Woodruff School of Mechanical Engineering, Georgia Institute of Technology, Atlanta, GA, USA; Department of Rehabilitation Sciences, Medical University of South Carolina, Charleston, SC, USA; Institute for Robotics and Intelligent Machines, Georgia Institute of Technology, Atlanta, GA, USA; Wallace H. Coulter Department of Biomedical Engineering, Georgia Institute of Technology & Emory University, Atlanta, GA, USA

**Keywords:** Motor Modules, Autoencoder, Muscle Synergy, Matrix Factorization

## Abstract

Motor module analysis is an important tool in the study of movement, particularly in people with impaired neural control. The most common method for computing motor modules is non-negative matrix factorization (NMF), which identifies a matrix of motor modules and their corresponding time-series activity from electromyography data. NMF has several limitations, including dependence of the muscle weightings on the number of modules selected. Approaches for selecting the number of modules vary between studies, making it difficult to compare and reproduce results. Some metrics of motor control complexity use the variance accounted for when extracting a single motor module (VAF_1_), yet that module’s structure offers little biomechanical interpretability. In this work, we present a method for computing motor modules using an autoencoder, a neural network architecture that can find latent representations of data. Using a single layer autoencoder, we extracted motor modules from data in able-bodied and individuals post-stroke. The structure of autoencoder-computed modules were significantly less sensitive to selected module number. With the autoencoder-computed modules, increasing the number of modules added new information, instead of splitting previous modules. Autoencoder-computed modules, especially at low module counts, had more distinct and interpretable biomechanical functions. Lastly, the autoencoder-computed modules are consistent with previous NMF studies in persons with stroke, which found fewer modules needed to explain the muscle activity of paretic limbs. Our autoencoder-based method offers a new approach for computing motor modules, with advantages of better stability in module structure across module counts, and a more biomechanically relevant interpretation of VAF_1_.

**NEW & NOTEWORTHY:** This work presents an approach for computing motor modules using an autoencoder and comprehensively compares the in stability of motor module structure, functional significance at low module counts, and interpretation of VAF_1_ to the current state of the art. The AE-computed module structures were more stable at different module counts. The AE has the potential to improve confidence in module structure and make analysis less dependent on the specific number of modules selected.

## INTRODUCTION

Motor module analysis has been a useful tool in understanding human motor control, particularly in the context of rehabilitation in populations with neurological injuries (1–5). Motor modules (also known as muscle synergies) are sets of coactivated muscles that describe structured spatial coordination of muscles by the nervous system to execute movements (1, 6, 7). These modules are computed by applying dimensionality reduction approaches to muscle activity data measured with electromyography (EMG). In analyses of individuals with neural control impairments, motor modules correspond with functional ability, and are particularly valuable in assessing impairment (2, 3, 8–11). For example, in gait, Clark et al. showed that individuals who have had a stroke typically had fewer modules compared to healthy individuals, and the number of modules was related to their functional ability. In studies comparing module structure between tasks, modules are partially preserved across tasks, suggesting motor modules represent a library of muscle patterns accessible by the central nervous system for movement (12–16). The relative weightings of the muscles within each module reflect biomechanically relevant coordination patterns used in the movements (1). In this way, knowing the number of modules and accurately determining their structure are important considerations for motor module analyses.

The current standard for computing motor modules, non-negative matrix factorization (NMF), has several challenges, specifically that the interpretation of motor module structures is highly dependent on the selected number of modules (17–19). NMF decomposes a matrix of EMG data into two non-negative matrices, which in the context of motor module analysis represent the groupings of muscles (module structure) and their activations over time (recruitment) (6). The number of modules is selected when executing the matrix decomposition algorithm, and module structure and recruitment is highly dependent on the number of modules selected. If the number of modules increases, the structures of the previous modules “splits”, making it difficult to determine the “true” module structure. For example, for an individual, one of the modules computed at a module count of two would split into two distinct ones at a module count of three. Thus, selecting two or three would lead to different conclusions about that individual’s motor control pattern available for movement. This problem is compounded by the fact that determining the number of modules remains an open problem. Typically, researchers iteratively perform NMF by adding modules until a threshold based on how much of the original EMG variance is accounted for by the module solution (VAF or R^2^) is reached. These thresholds are highly variable between studies (4, 20, 21), making comparison between studies difficult. Additionally, it can be difficult to compare and reproduce across studies because selecting the number of modules can be highly dependent on various factors, such as filtering level, averaging, or the presence of noise (18). Often, these parameters are heuristically selected based on experience. The lack of consistency in module structure across module counts means that changes in how module counts are selected can meaningfully change conclusions from motor module analysis.

One method often used to address this variability with module analysis methods is to use the variance accounted for by a single module (VAF_1_) for quantifying motor control complexity (22–24). This approach does not assume that only a single module exists at the level of the nervous system, rather it quantifies motor control based on the idea that more variability accounted for by a single module reflects a less complex control strategy. VAF_1_, or similarly derived metrics such as Walk-DMC, have strong relationships between functional ability in children with cerebral palsy, and correlate well with clinical measures (22, 25–27). Its ease of use, along with the intuition behind the approach, makes using a single module an attractive option for researchers aiming to quantify the motor control complexity of their participants. However, the resultant module computed with NMF does not group muscles into modules that reflect coordination patterns to produce distinct biomechanical functions, as it often contains every muscle that was part of the collection (1, 6). When analyzing how motor modules change during or after some sort of intervention, single module metrics do not allow for interpretation of specific muscle coordination changes and cannot isolate changes in module recruitment. Therefore, even if there are changes in VAF_1_, this may not reflect changes to the biomechanical function of a *specific* module. An improvement to the biomechanical interpretability of VAF_1_ could serve to make single module analysis more informative.

Autoencoders are an approach to motor module computations that can alleviate some of the issues typically associated with the current methodological approaches using NMF. Autoencoders are a dimensionality reduction technique that utilizes a neural network structure to encode latent representations of data (28). In principle, autoencoders are similar in function to other dimensionality reduction approaches, such as PCA, ICA, or NMF, with a few added advantages. Autoencoders function by using a symmetrical neural network, with an encoder on one side, a decoder on the other, and a “bottleneck layer” (latent space) between them. The size of the latent space is smaller than the input/output, forcing the model to learn a lower dimensional encoding of the data. Studies that use an autoencoder to compute motor modules found similar reconstruction accuracy from the AE-based method as NMF (29–31), however, the module structures were not compared.

In this work we used the most standard autoencoder implementation to extract motor modules. Here, we perform a comprehensive analysis comparing the structure and stability of motor modules computed with an Autoencoder (AE) to modules computed with NMF. Here we show that an autoencoder can produce more consistent module structures across module counts and improve interpretability of module biomechanical function. We evaluated how motor module structures computed by an autoencoder compare to those computed with NMF, and how stable these structures are across module counts. Additionally, we evaluated how the number of modules selected differs between the two methods. We also showed how the autoencoder implementation impacts analysis of metrics such as VAF_1_ in the context of overall motor complexity and changes to biomechanical function. We applied our autoencoder extraction technique to a previously published dataset of stroke subjects to demonstrate its utility in the analysis of individuals with altered motor control ability (2). We evaluated whether prior conclusions, such as fewer motor modules, or a higher VAF_1_ in the paretic limb, were replicated when modules were computed using the AE.

## MATERIALS AND METHODS

Electromyography (EMG) from two datasets were used to evaluate the autoencoder and NMF-based methods. The first dataset (Camargo et al. 2021) consisted of 11 muscles collected from the right leg from 21 healthy able-body individuals, with all persons walking at 1.3 m/s (32). The muscles collected for the healthy EMG data were the medial gastrocnemius (MG), tibialis anterior (TA), soleus (SOL), vastus medialis (VM), vastus lateralis (VL), rectus femoris (RF), biceps femoris (BF), semitendinosus (SEMT), gracilis (GRA), gluteus medius (GM), and external oblique (EXOB). The second dataset from Clark et al. 2010 consisted of 8 muscles collected from the paretic and non-paretic legs of 52 participants post-stroke walking at self-selected walking speed (2). The muscles collected for the post-stroke data were the gluteus medius (GM), lateral hamstring (LH), medial gastrocnemius (MG), medial hamstring (MH), rectus femoris (RF), soleus (SOL), tibialis anterior (TA), and vastus medialis (VM). All EMG was bandpass filtered between 20 and 400 Hz, demeaned and rectified, and low-pass filtered at 10 Hz. Each participant was evaluated as a single, 30-second trial of continuous walking data. For participants post-stroke, modules were computed independently for the paretic and non-paretic legs. EMG data was pre-processed in MATLAB. Motor modules were computed in Python, with the `non_negative_factorization` function from the scikit-learn package, and an autoencoder built using tensorflow/keras packages, for the NMF and AE approach, respectively(33, 34). Additional details regarding sample demographics, inclusion and exclusion criteria, and study designs can be found in their respective publications.

We used a single hidden-layer autoencoder, where the input data is directly connected to the latent space, and the latent space is directly connected to the output, by fully connected neural network layers (Figure 1A) (28). The input and output shapes were equal to the number of muscles collected, and the size of the latent space represented the number of motor modules. At each layer, a rectified linear unit (ReLU) activation function was applied, and the layer weights were constrained to be non-negative. ReLU constrains the layer outputs to zero if they would be less than zero. A constant bias term is also added at each layer. Figure 1B describes the parameterization for both the NMF and AE models for motor modules. The AE-computed motor modules were extracted by taking the trained weights between the latent space and output (decoder). The time-series recruitment of each module was the activity of the corresponding dimension of the latent space for a given input.

**Figure 1.**
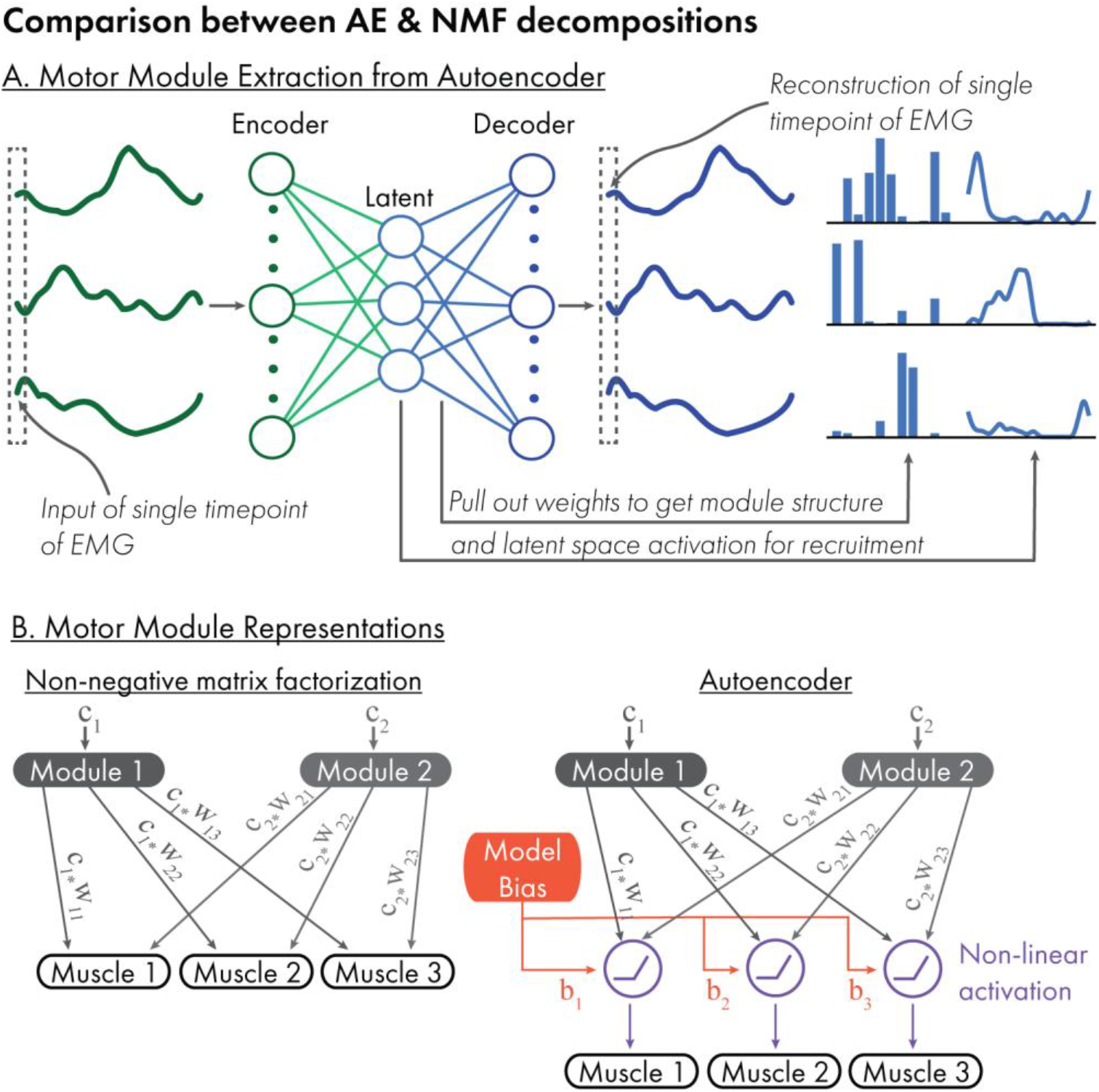
A) Layout of utilized autoencoder structure, and how motor module structures and activation profiles were computed from the trained model. The size of the latent space (number of nodes in the middle layer) determines the number of modules. B) Model for reconstruction using the NMF (left) and AE (right) based methods. The AE includes two additional terms, a constant bias and a non-linear activation function (ReLU), which allow for a more complex parameterization of the relationship between module activation and muscle activity.

We evaluated the similarity of the motor module structure of corresponding modules as we incremented the module count in both the AE and the NMF computed modules. For each method, modules were computed iteratively, starting at 1 module and progressing up to 8. We defined agreement in module structure at differing module counts as the similarity between each module and the most similar module at the previous module count, measured by Pearson’s correlation. A higher agreement indicates that the module structure is better preserved across module counts, and that the motor modules exhibit less “splitting” as the number of modules is incremented. A high agreement suggests the information contained within a module is approximately equivalent, regardless of the number of modules selected. To quantify the biomechanical functional significance of each module, we computed the modulation index for each module activation over the gait cycle, which aims to capture how distinct the largest points of activation are from the smallest points of activation (35). Modulation index (MI) was defined as

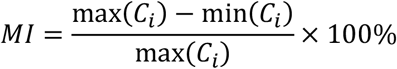

where *C*_*i*_ is a vector of the activation of a given motor module *i* over the gait cycle. Differences between the NMF and AE method at each module count were statistically examined with paired t-tests.

To evaluate the reconstruction quality, we computed VAF for both methods. We computed VAF at each module count and additionally looked at VAF_1_ specifically. We also wanted to evaluate how much the module structure alone contributed to the reconstruction, without the contribution of the autoencoder bias term. To do this, we manually set the bias component of the model to zeros and performed the same reconstruction and VAF computation. For a module count of one specifically, we examined how physiologically interpretable the module activation and structures were, to determine the impacts on VAF_1_ interpretation. We compared how distinct the activation profile was, and which muscles were included in the single module produced by the two methods.

We also tested whether the number of modules computed differed between the two methods (36). We computed the reconstruction accuracy with a bootstrapped approach. Each set of EMG input had 80% of its data randomly selected and evaluated against its corresponding reconstruction. This was repeated 100 times and for each module count. When the 95^th^ percentile of the bootstrapped VAFs exceeded 90%, that number of modules was selected (6). Lastly, we looked at the VAF at each module count to evaluate if there were systematic differences in the reconstruction accuracy as modules were added.

To demonstrate utility of the AE computed motor modules in a population with motor control deficits, we repeated the methods using the dataset from Clark et. al. Additionally, we computed the relative distribution of VAF at each module count between the paretic and nonparetic limbs. This allowed us to see whether typical trends in VAF at each module count reported in previous work were still observed when modules were computed with the AE. Motor modules were independently computed for the paretic and nonparetic legs.

## RESULTS

### Able-Bodied Individuals

At a module count of one, the AE-computed modules computed from the healthy individuals’ EMG data had a more distinct activation than the NMF-computed modules, which can allow for a more biomechanically relevant interpretation of VAF_1_ and other similarly derived metrics. Where the NMF-computed first module included essentially all the muscles at a similar magnitude, the AE-computed first module had a distinct structure consisting primarily of knee extensor muscles, (VM, VL, RF) along with the GM, that was preserved as modules were added (Figure 2. A/B, row 1). Further, the NMF-computed first module was on throughout the gait cycle, whereas the AE-computed module was on primarily in stance phase and off during swing phase (Figure 2. A/B, row 1). Accordingly, the AE-computed first module had a significantly higher modulation index than the NMF-computed module (98.3% ± 1.3% and 62.1% ± 12.7%), Indicating more phase-dependent activation of the module throughout the gait cycle. When looking at VAF_1_ for both methods, the total reconstruction is similar (VAF=73.7% ± 6.4% and 68.2% ± 7.5%, for the AE and NMF, respectively), with slightly higher reconstruction for the AE (Figure 3B). When the bias terms were zeroed out, representing only the component of the data accounted for by the first module, the AE-computed first module had a significantly lower VAF_1_ (35.1% ± 8.0%) than the NMF approach.

**Figure 2.**
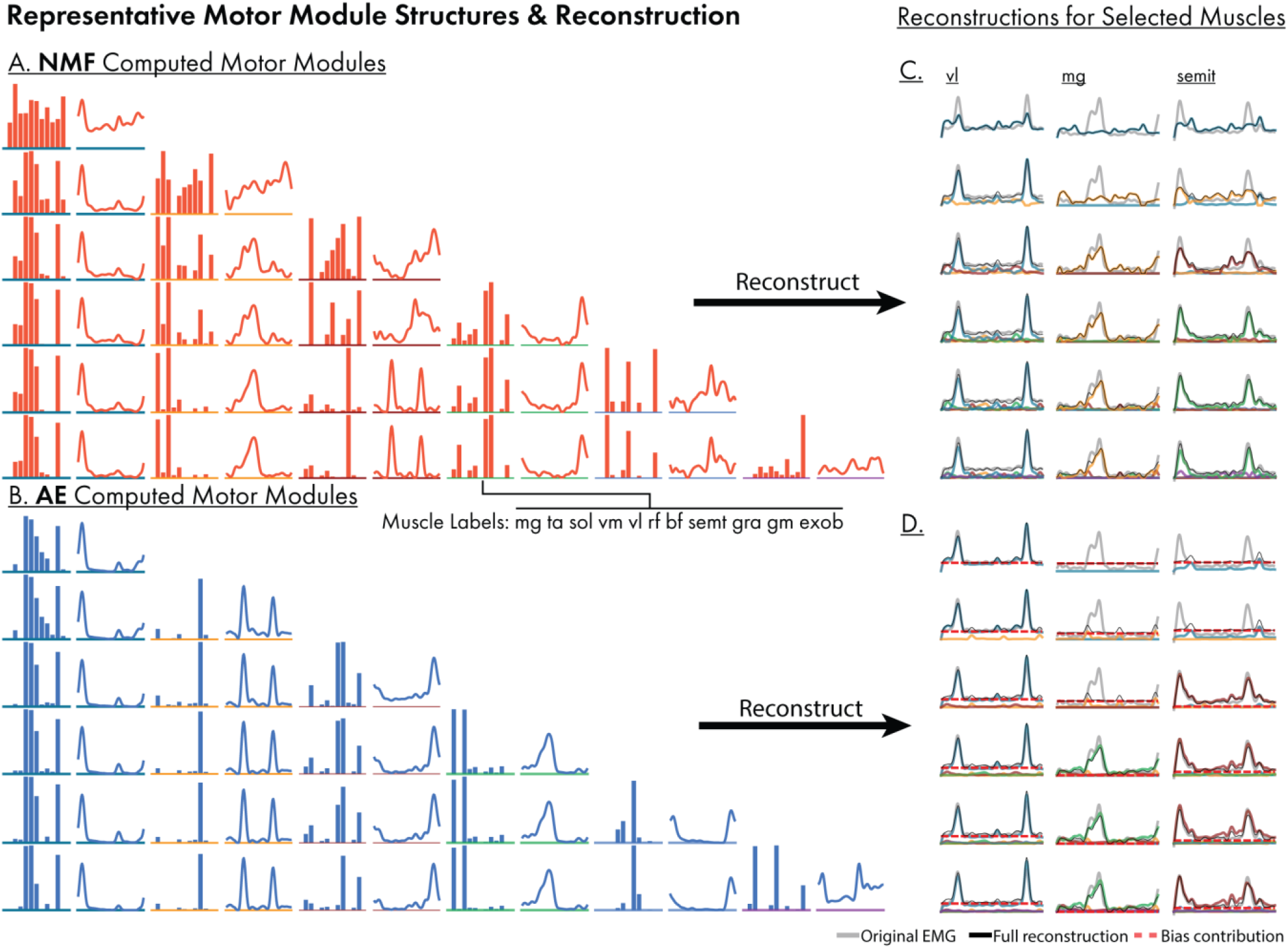
A-B) A representative set of modules computed at differing module counts for the NMF and AE method respectively. The AE shows better retained module structures as the number of modules is incremented. * indicates a significant difference between the AE and NMF based methods. C) Reconstruction comparison between the NMF and autoencoder methods, for the medial gastrocnemius, vastus lateralis, and semitendinosus muscles. The colored traces show the contribution of each module, corresponding with the same-colored module in A. The autoencoder method shows a clearer fitting at higher modules counts, with the majority of reconstruction at

**Figure 3.**
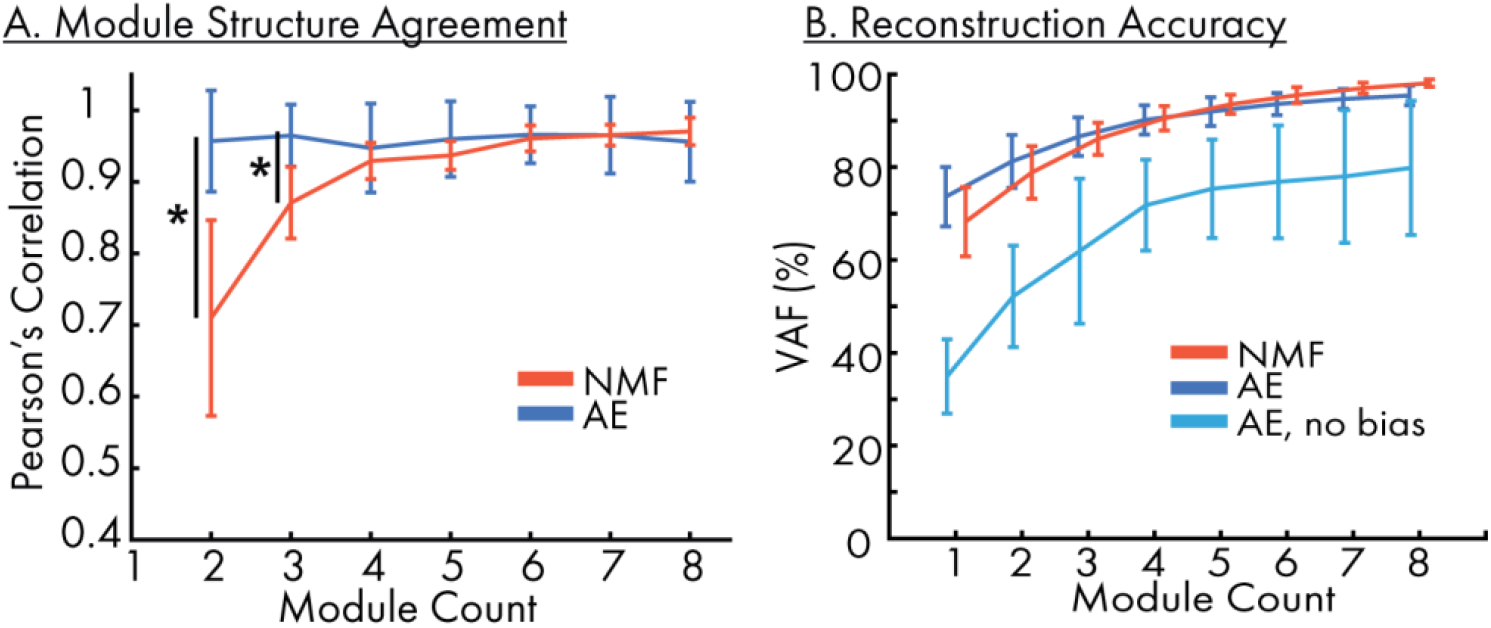
A) The agreement in module structure as module count is incremented for all subjects as measured by Pearson’s correlation. Error bars indicate one standard deviation. B) Overall reconstruction accuracy from the two methods. The autoencoder shows a slightly higher VAF1 due to the extra parameterization of the bias term.

There was no significant difference between the two approaches in the number of modules selected by AE vs NMF for the EMG data from healthy individuals. This suggests that overall, the number of modules determined for an individual are likely to be similar whether NMF or an AE is used to extract the modules. However, the AE-module reconstructions using zeroed bias terms had a lower VAF, like with VAF_1_, with this difference reducing as the module count increases (Figure 3B). In this case, the bias term may represent some sort of error component where the information not relevant to the module structure are part of the bias term, which shrinks as modules are added. The MI for the subsequently computed modules were significantly higher in the AE-computed modules compared to the NMF-computed modules up to a module count of 8, though with a much closer value at a module count of 3 or higher, indicating that the improved distinction in biomechanical function of the first module is still partially present when computing higher numbers of modules.

The AE-computed module structures from the healthy individual EMG data were much more stable across modules counts, making module analysis results much less sensitive to the number of modules selected. The AE-computed modules had a stronger agreement in module structure and less “splitting” as the module count was incremented, when compared to the NMF-computed modules. In contrast to the AE-computed modules, the NMF-computed modules had substantially different structures within each module depending on the selected number of modules. The NMF module structures included all muscles when one or two modules were computed (Figure 2A, first row). As module count increased, the NMF-computed modules split into more distinct ones (Figure 2A). Conversely, the biomechanically distinct structure of AE-computed module at a module count of one (Figure 2B, first row) was retained as the module count increased. Pearson’s correlation coefficient between the most similar modules at each module count was significantly higher in the AE than NMF for module counts of 2 and 3 (Figure 3A). The AE-computed module stability is also demonstrated in the EMG reconstructions (Figure 2C/D). In the representative example at one module, NMF partially reconstructs the VL, and the AE method fully fits the VL. The MG and SEMIT EMGs are partially fit using 1, 2, and 3 NMF modules, but within different modules depending on how many total modules are used. In the NMF reconstructions, muscle weightings and features of the activation present in one module move to the others as more modules are added. In the AE case however, MG and SEMIT are essentially unfit at modules counts 1 and 2, with most of the reconstruction due solely to the bias term. At 3 modules, the SEMIT is properly reconstructed as the new module contains its relevant information, and then the same happens to the MG at 4 modules.

### Individuals Post-Stroke

Like in the able-bodied dataset, the AE results in more biomechanically distinct first modules than the NMF method, comparing both the structures visually and the modulation index (90.5% ± 11.0% and 56.7% ± 14.13% for the AE and NMF-computed modules, respectively) (Figure 4A/B). Accordingly, in both the paretic and non-paretic limbs, the agreement of modules structure across module count remained significantly higher in the AE-computed modules than in the NMF-computed modules.

When the AE-reconstruction for the post-stroke individuals was computed with a zeroed out bias term, there was no significant difference in reconstruction quality between the paretic and non-paretic limbs at any given module count (as measured by VAF) and only showed a nominal higher paretic side VAF at a module count of one (Figure 4C, dotted green). However, when computing the contribution to the reconstruction of only the bias term (motor module weights zeroed), the paretic side limb had a higher VAF (significant at module counts 2-6), potentially indicating more tonic activity (Figure 4C, dashed purple). When evaluating using the full reconstruction (i.e. bias included), the number of selected modules from both the AE and NMF approaches was lower in the paretic limb than in the non-paretic limb, aligning with expected results when evaluating the motor complexity of post-stroke individuals (Figure 4D). Although the relative number of modules between the paretic and non-paretic limbs were similar between the two methods, the AE approach did tend to identify some individuals requiring more selected modules across both the paretic and non-paretic limbs, though this was not a systematic increase in all participants.

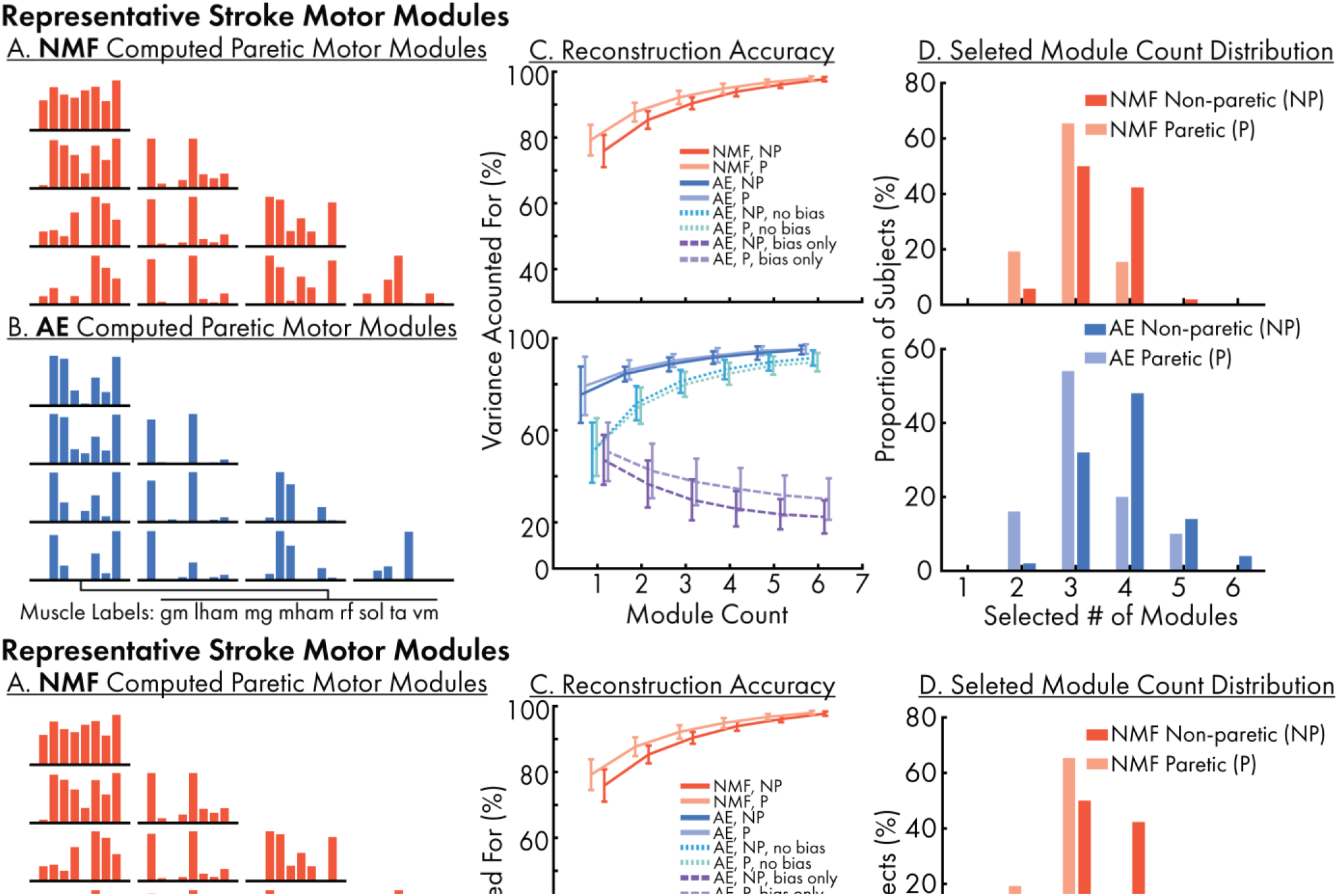

## DISCUSSION

Here we demonstrate the advantages of using an autoencoder to compute motor modules, or muscle synergies, from locomotor electromyography data during walking. We evaluated the characteristics of module structures and activations from both methods and demonstrated that the AE-computed module structure and activations were more stable at different module counts than the NMF-computed modules. The AE-computed modules also improved the biomechanical interpretability of the first module and analysis of VAF_1_. The autoencoder showed these advantages in both the healthy EMG data, and in EMG data from individuals post-stroke, allowing for more consistent motor module analysis in a clinical population.

Computing a single module using the autoencoder captures how much a *specific* motor module contributes to the total muscle activity, regardless of the number of modules extracted. Typically, the first AE-increased for paretic and non-paretic limbs in individuals who have had strokes, computed for NMF-modules, and AE-modules (with reconstructions including the bias and only using the bias). Paretic side data has significantly higher reconstrution accuracy thannon-paretic side data at each module count. Right) Distribution of selected number of modules when using the standard approach for paretic and non-paretic limbs in individuals who have had strokes. The non-paretic data for both the AE and NMF methods show a distribution further to the right and has a statistically significant increase in number of selected modules when compared to the paretic data.

motor module consists of an extensor module that is active during stance phase. The AE-based module therefore eliminates the issues of having to select the “correct” number of modules using NMF to make comparison across individuals or interventions. Further, extracting a single AE-motor module reflects physiologically-modulated biomechanical coordination pattern, whereas a single NMF-motor module, often used as a proxy for motor control complexity (22, 26, 37, 38), reflect more of the mean activation of muscles. The addition of a bias term when using the autoencoder likely acts as an error term to account for muscle activations that are not well-described by the number of extracted modules. Therefore, zeroing out the bias term enables the assessment of the identified motor modules, even if there are additional motor modules contributing to the motor pattern. Ultimately, the two methods can produce similar results in terms of the number of modules selected and final motor module structure accounting for most of the variability in the data.

The inclusion of the bias term in the autoencoder parameterization of the motor modules likely produces some level of regularization, preventing the modules from “splitting” when the module count is increased. Because the NMF model’s parameterization is limited to the weights and activations of each module, the fitting process must account for as much data as possible with the limited terms, to minimize the Frobenius norm between the data and the reconstruction (39, 40). In contrast, the AE decoder is modeled by the activation, weights, and an additional bias term applied to all muscle activity. In the single module case, the bias can be conceptualized as pseudo-module with constant activation, which may have little meaning in terms of biomechanical function, but prevent the first module from representing the mean activity across all muscles. Importantly, the bias allows for the neural network training to more selectively tune the weights between the latent space and the output such that they represent consistent underlying motor patterns and placing information unrelated to the module(s) in the bias term(s). Therefore, the total VAF of the AE reconstruction is higher than NMF when including the bias, but lower than NMF when the bias is zeroed out and reflects only the variability accounted for by the identified modules. In this sense the bias can be considered as an error term not part of the phasic motor modules. The inclusion of the bias term in the AE model also has the potential to generate more plausible modules when muscles with very low or tonic activity are included in the analysis.

The AE-computed modules have properties that are beneficial for motor module analyses for comparisons within and between individuals. In the NMF case, because the modules show a “splitting” behavior, where one module’s information splits off into two others as the module count increases, it can be unclear which muscles are grouped together, and what is simply an artifact of the NMF algorithm (6). Additionally, there is no consistent method of determining the number of motor modules, and with the NMF-computed modules, this can have a significant impact on subsequent analysis (18, 19, 41–44). Because the structures of the modules produced by the AE are relatively well preserved across module counts, the overall analysis is less dependent on selecting the “true” module count, and researchers can have more confidence in their analysis without having to arbitrarily select thresholds or a fixed number of modules (2, 45). The consistency of motor modules thus allows the extracted motor modules to be interpreted functionally and represents an individual’s consistent muscle coordination patterns underlying the observed EMG patterns regardless of the number of motor modules extracted. Therefore, changes in the structure of motor modules can be more effectively assessed across tasks, before and after intervention, and between individuals (17, 42, 46, 47).

The AE-computed modules gave similar results in post-stroke individuals as the NMF approach, consistent with results from prior literature, and could be used in place of the typical methods without significantly changing expected outcomes, while improving confidence in module structure. Previous research has reported fewer motor modules that account for a high VAF in the paretic limbs of individuals who have had strokes (2). We demonstrate that this is still present when these metrics are computed from the AE-based modules. Notably, the higher expected paretic side VAF is not present when we perform the reconstruction without the bias term and only returns when computing with the bias term. This further suggests that the bias term is generally capturing either tonic activation, or highly variable features stride to stride, which are best represented by an invariant average, which is expected to be more present on the paretic side limb. Additionally, the benefits of the AE approach are preserved in the impaired case, which can improve the overall analysis by giving more confidence in module structure at different module counts. This can be even more impactful in the case of stroke data, as not only are fewer modules required, but the impaired motor modules can be shown to be merged forms of healthy modules (2, 3, 42). However, with NMF-computed modules, it is not necessarily clear if this merging is a behavior of the nervous system, or a byproduct of the algorithm and module count selection. Using an AE to compute motor modules can give more consistent modules, where merged modules are more likely to be physiologically rooted, as opposed to being a byproduct of too few modules extracted. It should be noted that our results do not exactly reflect the results shown in Clark et. al, as the exact filtering process differed, and we computed our modules using full time series EMG data, not gait cycle-averaged data.

It remains to be seen whether AE-computed modules will have improved stability in other cases where NMF falls short. For example, when a muscle is minimally modulated during a task (either from tonic activity or technical data collection problems), the bias term may be able to better capture that instead of splitting the muscle up between modules it isn’t part of. Another potential instance where the AE may perform well is in instances where a muscles contribution to a module is non-linear. The ReLU activation function, for example, can make a negative bias term act like a threshold, meaning that the module must be activated above a certain level for some muscles to start modulating. Other activation functions may be better suited to different tasks. It also has yet to be shown how the AE-computed modules behave in the presence of noise. For instances of constant noise, it may have a similar behavior to the tonic activation, where the bias term accounts for a baseline level of activation. However, in the presence of large motion artifacts, it may have the issue of allocating a whole motor module to that apparent activation.

## APPENDIX

## DATA AVAILABILITY

Code used to compute motor modules using the described method can be found at https://github.com/Neuromechanics-Lab/motor-module-autoencoder.

Source data for this study from Camargo et al., 2021 are openly available in three parts at https://data.mendeley.com/datasets/fcgm3chfff/1, https://data.mendeley.com/datasets/k9kvm5tn3f/1, and https://data.mendeley.com/datasets/jj3r5f9pnf/1.

Source data for this study from Clark et al., 2010 are available from the NIH Center of Biomedical Research Excellence in Stroke Recovery upon reasonable request. Please contact the authors to inquire about access to the data.

## ACKNOWLEDGEMENTS

We would like to thank Dr. Michael Rosenberg for his feedback on this project and the data processing. This work was supported by the National Science Foundation Graduate Research Fellowship Program (NSF GRFP DGE-2039655) and the Georgia Institute of Technology George W. Woodruff School of Mechanical Engineering Interdisciplinary Research Fellowship.

